# Establishment of an orthodontic retention mouse model and the effect of anti-c-Fms antibody on orthodontic relapse

**DOI:** 10.1101/575589

**Authors:** Jiawei Qi, Hideki Kitaura, Wei-Ren Shen, Akiko Kishikawa, Saika Ogawa, Fumitoshi Ohori, Takahiro Noguchi, Aseel Marahleh, Yasuhiko Nara, Itaru Mizoguchi

**Affiliations:** Division of Orthodontics and Dentofacial Orthopedics, Department of Translational Medicine, Tohoku University Graduate School of Dentistry, 4-1 Seiryo-machi, Aoba-ku, Sendai 980-8575, Japan

**Keywords:** orthodontic tooth movement, relapse, osteoclast, anti-c-Fms antibody, retention

## Abstract

Orthodontic relapse after orthodontic treatment is a major clinical issue in the dental field. However, the biological mechanism of orthodontic relapse is still unclear. This study aimed to establish a mouse model of orthodontic retention to examine how retention affects the rate and the amount of orthodontic relapse. We also sought to examine the role of osteoclastogenesis in relapse using an antibody to block the activity of M-CSF, an essential factor of osteoclast formation. Mice were treated with a nickel-titanium closed-coil spring that was fixed between the upper incisors and the upper-left first molar to move the first molar in a mesial direction over 12 days. Mice were randomly divided into three groups: group 1, no retention (G1); group 2, retention for 2 weeks (G2); and group 3, retention for 4 weeks (G3). In G2 and G3, a light-cured resin was placed in the space between the first and second molars as a model of retention. Orthodontic relapse was assessed by measuring changes in the dimensions of the gap created between the first and second molars. To assess the activity and role of osteoclasts, mice in G3 were injected with anti-c-Fms antibody or PBS, and assessed for changes in relapse distance and rate. Overall, we found that a longer retention period was associated with a slower rate of relapse and a shorter overall amount of relapse. In addition, inhibiting osteoclast formation using the anti-c-Fms antibody also reduced orthodontic relapse. These results suggest that M-CSF and/or its receptor could be potential therapeutic targets in the prevention and treatment of orthodontic relapse.

## Introduction

Orthodontic relapse following orthodontic treatment has been a major clinical issue for orthodontic dentists and patients. Retainers, which are the most widely used device in the clinical setting, must be worn for at least a few years to prevent relapse after the completion of orthodontic treatment. In earlier work, it was advocated that permanent retention may be the only solution to maintain a long-term post-treatment effect [1]. Yet, in some cases, teeth begin to relapse to their original position even after orthodontic retention. It has been suggested that a relapse force is generated during orthodontic tooth movement and stored in the periodontal and transseptal fiber systems [2]. After the orthodontic appliance is removed, the relapse force is released, and the teeth begin to move back to their original positions [3]. Indeed, there is more than a 19% relapse rate even with the effective use of retainers after orthodontic treatment at 3 years [4]. An orthodontic retention animal model is necessary to elucidate the mechanism of orthodontic relapse. However, as yet, there is no animal model of retention with which to evaluate these processes.

Orthodontic tooth movement (OTM) is achieved by continuous alveolar bone resorption by osteoclasts on the side undergoing compression along with the stimulation of new bone formation by osteoblasts on the side subjected to tension. Osteoclasts, derived from bone marrow cells, regulate bone resorption during bone remodeling. Studies of orthodontic relapse show that there is a tendency for there to be higher numbers of osteoclasts in association with a greater distance of tooth movement [5-7]. Osteoclast differentiation is dependent on macrophage colony-stimulating factor (M-CSF) and the ligand for the receptor activator of necrosis factor κB (RANKL) [8]. Tumor necrosis factor (TNF)-α is also essential for osteoclast induction [9-11]. In previous studies, we showed that an antibody against the receptor for M-CSF, c-Fms, can inhibit TNF-α–induced osteoclast formation in vitro [6] and in vivo [12]; LPS-induced osteoclast formation [13]; and arthritis-induced osteoclast formation [12]. Delivery of an anti-c-Fms antibody can also inhibit orthodontic tooth movement, by blocking osteoclastogenesis and bone resorption [6], and inhibit root resorption during orthodontic tooth movement [14].

Several studies have reported that tooth movement might be controlled by pharmacological therapy, with tooth movement able to be suppressed by the administration of bisphosphonates [15, 16] and osteoprotegerin [17] in animal models. Likewise, simvastatin [18], relaxin [19], low-level laser therapy [20] and aspirin [21] have been associated with preventing orthodontic relapse in animal studies. The degree of relapse can be controlled by modifying the dental supporting tissues. However, the mechanism of orthodontic relapse is still unknown and there has been no study of orthodontic relapse after retention. A greater understanding of the relapse process is required to determine ways to inhibit relapse or reinforce retention.

In this study, we established a mouse model of orthodontic retention. Using these mice, we investigated the relapse distance, relapse rate, and level of osteoclast activity after orthodontic tooth movement with or without retention, and evaluated the effect of anti-c-Fms antibody treatment on orthodontic relapse using a mouse model.

## Materials and Methods

### Experimental Animals

C57BL6/J mice at 10 to 12 weeks were obtained from CLEA Japan Inc. (Tokyo, Japan). The mice were fed in cages in a room maintained at 21–24°C with a 12-hour/12-hour light/dark cycle. The granular diet (Oriental Yeast, Tokyo, Japan) was provided to prevent eating difficulty during force-loading. Mice were anaesthetized in each experiment. A combination anesthetic including medetomidine, midazolam and butorphanol was administered to mice by intraperitoneal injection. In order to minimize the suffering, mice were euthanized by inhalation of an overdose of 5% isoflurane. All experimental procedures conformed to “Regulations for Animal Experiments And Related Activities at Tohoku University”, and were reviewed by the Institutional Laboratory Animal Care and Use Committee of Tohoku University, and finally approved by the President of University.

### Orthodontic tooth movement

Mice were fit with an orthodontic appliance, as described previously [6, 7]. Briefly, under anesthesia, a nickel titanium closed coil spring (Tomy; Fukushima, Japan) was fixed between the upper incisors and the upper-left first molar. A 0.1-mm stainless steel wire was then used to move the first molar in a mesial direction. According to the database of the manufacturer, the force level of the appliance after activation is approximately 10 g (Fig. 1A). Orthodontic tooth movement (OTM) was achieved after forced loading for 12 days. The distance between first molar (M1) and second molar (M2) was measured. A tray containing hydrophilic vinylpolysiloxane (EXAFAST Injection Type, GC Co., Tokyo, Japan) was placed onto the maxillary teeth to obtain an impression. The distance between the distal marginal ridge of the first molar and the mesial marginal ridge of the second molar (dotted line) was measured to assess tooth movement (red double arrow) by stereoscopic microscopy (VH-7000; Keyence, Osaka, Japan) (Zaki et al., 2015). Space retention was considered successful when movement was less than 10 μm of the original OTM (Fig. 1B).

**Fig. 1.**
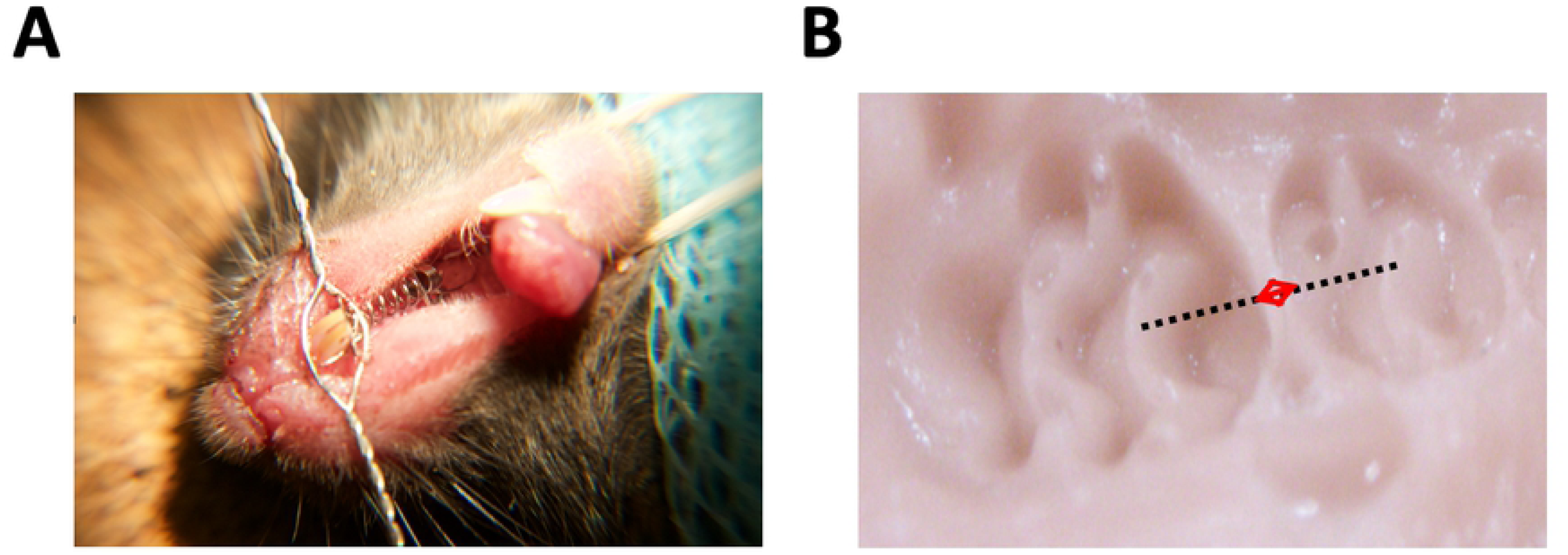
Mouse model of orthodontic retention. (A) A nickel-titanium closed-coil spring is fixed between the upper incisors and the upper-left first molar. A 0.1-mm stainless steel wire is used to move the first molar in a mesial direction. (B) Photograph of the silicone impression after tooth movement. The dashed line connecting the central fossae of the first and second molars was used to measure the distance of tooth movement (from the distal marginal ridge of M1 to the mesial marginal ridge of M2) (red double arrow).

### Mouse model of orthodontic retention

After OTM for 12 days, the appliances were removed, and mice were randomly divided into three groups: No retention (Group 1, G1); Retention for 2 weeks (G2); and Retention for 4 weeks (G3). For mice in G1, the period of relapse (r0) began immediately after the appliances were removed. For mice in G2 and G3, resin was placed within the created gap for 2 weeks or 4 weeks. The period of relapse began after removal of the appliance (G1) or resin (G2 and G3). There were eight mice in each group. Four mice were sacrificed on day 0 (r0). The other four mice were anaesthetized and used to measure the retention of space during the orthodontic relapse period every day for the first 5 days and then every second day until day 15 (see Fig. 2A, red dots). These mice were sacrificed on day 15 (r15) in each group.

**Fig. 2.**
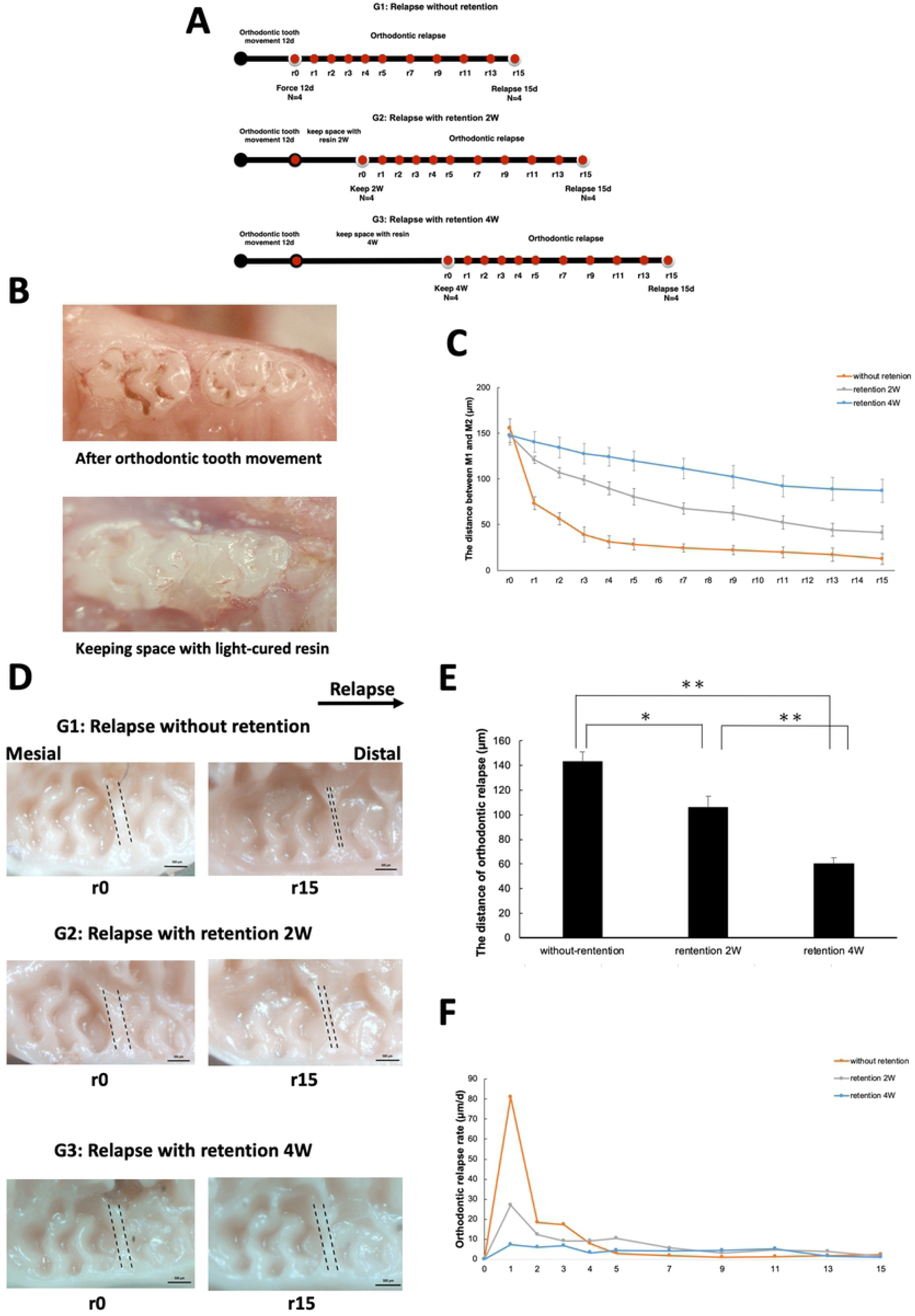
Evaluation of relapse distance and rate with or without retention. (A) Experimental timeline for the establishment of the mouse retention model. Mice were randomly divided into three groups: No retention (Group 1, G1); Retention for 2 weeks (G2); and Retention for 4 weeks (G3). There were 8 mice in each group. All mice received orthodontic tooth movement for 12 days. For mice in G1, the period of relapse began immediately after the appliances were removed. For mice in G2 and G3, resin was placed within the created gap for 2 weeks or 4 weeks. The period of relapse began after removal of the resin (G2, G3) or orthodontic appliance (G1). Orthodontic relapse was measured on the days indicated by the red markers (r0, r1, r2, etc). Four mice were sacrificed on day 0 (r0) and another four on day 15 (r15) in each group. (B) Photograph of the light-cured resin used as a retainer to maintain space between the first molar (M1) and second molar (M2). (C) Changes in the distance of orthodontic relapse in each group (measured in µm. *n* = 4 for each group. (D) Relapse distances were measured by taking silicone impressions. The arrow (top) represents the direction of relapse. Scale bars = 500 μm. (E) Comparison of relapse among the three groups. The relapse distance decreased with an increasing retention period. *n* = 4 for each group. \**P* < 0.05; \*\**P* < 0.01. (F) Relapse rate was rapid initially in all groups, and then gradually decreased toward the end of the experimental period. *n =* 4 for each group.

For mice in the retention groups (G2 and G3), a light-cured resin (GC Co., Tokyo, Japan) was used to maintain the space created between M1 and M2. Under anesthesia, the OTM appliances were removed, and a dental etching agent (GC Co., Tokyo, Japan) was smeared onto the tooth surface to create a large adhesive area. The tooth was then washed with water and dried with medical cotton. A dental bonding agent (GC Co., Tokyo, Japan) was used to fix the resin after LED light activation for 5 sec. Light-cured resin was placed into the interdental space between M1 and M2, and solidified with LED light for 20 sec (Fig. 2B).

### Histological preparation and analysis

After sacrifice (r0 and r15), the maxillae were removed and fixed in 4% paraformaldehyde (PFA) for 24 h at room temperature. The tissue was demineralized in 14% ethylene diamine tetra-acetate (EDTA) for 3 weeks at room temperature, dehydrated in ascending concentrations of ethanol, embedded in paraffin, and sectioned at 4 μm for histological analysis.

Horizontal sections of the distobuccal region from the first molar bifurcation area to the apical root were prepared. Five levels from bifurcation area were evaluated in each sample: 100, 140, 180, 220, and 260 μm. The sections were deparaffinized and stained with tartrate-resistant acid phosphatase (TRAP) and hematoxylin. Naphthol-ASMX-phosphate (Sigma-Aldrich; St Louis, Missouri, USA), Fast Red Violet LB Salt (Sigma-Aldrich), and 50 mM sodium tartrate were used for TRAP staining. The sections were evaluated using light microscopy. Osteoclasts were counted on the mesial and distal sides. Cells were considered to be osteoclasts if they were TRAP-positive, multinucleated, and are located on the bone surface.

### Treatment with Anti-c-Fms Antibody

AFS98 is a rat monoclonal anti-murine c-Fms antibody (IgG2a) that inhibits M-CSF-dependent osteoclast formation by blocking the binding of M-CSF to its receptor [22]. An AFS98 hybridoma was cultured in HyQ-CCM1 medium (Hyclone, Logan, UT, USA), and the antibody was purified with Protein G (Sigma-Aldrich). Mice were injected every 2 days for a total of 9 injections with 1.5 μg of the anti-c-Fms antibody in 3 μL of phosphate-buffered saline (PBS). PBS only was used as a vehicle control. Injections were directed into the palatal gingiva close to the space between upper-left first and second molars during orthodontic relapse with a micro-injection apparatus (Hamilton, Nevada, USA).

### Statistical analysis

All data are presented as the mean ± standard deviation (SD), and were assessed by Scheffe’s F-tests and Student’s *t*-tests.

## Results

### Orthodontic tooth movement and orthodontic relapse

The timeline in Figure 2A shows the experimental design for the establishment of the orthodontic retention mouse model, including the timing of orthodontic tooth movement, the space retainment with resin, and the period of orthodontic relapse. Tooth movement in the mesial direction was observed for all mice after forced loading for 12 days. The mean distance between the upper left M1 and M2 was 150.19 ± 9.62 μm, with some individual differences seen. In all of the experimental groups, there was a decrease in the distance between M1 and M2 on day 15 of relapse (r15) as compared with day 0 of relapse (r0). These findings indicated that mice in all of the groups suffered orthodontic relapse, irrespective of retention (Fig. 2C). The best space retention at r15 was for mice in G3 (87.21 ± 12.59 μm) as compared with mice in G2 (41.46 ± 7.26 μm) and those in G1 (12.48 ± 6.04 μm). The interdental distance after 2 weeks was significantly lower in the control group on r15 as compared with the mice subjected to retention for 2 weeks (*P* < 0.01). Greater interdental distance was seen for mice subjected to retention for 4 weeks compared with those in the 2-week group (*P* < 0.01; Fig. 2C and 2D).

### Orthodontic relapse is inhibited by retention

The relapse distance was calculated as: relapse distance at r0 – relapse distance at r15. Relapse distance was significantly shorter (60.44 ± 4.91 μm) in the mice subjected to 4 weeks of retention (G3) as compared with those subjected to only 2 weeks of retention (G2, 106.18 ± 8.68 μm) or no retention (G1, 143.17 ± 7.83 μm) (Fig. 2E).

### Relapse rate is reduced after retention

Some relapse occurred in all groups, irrespective of treatment. The relapse rate peaked on day 1 among mice in G1 as compared with mice in G2 and G3. After day 1, the relapse rate declined, but remained higher for mice in G1 than for those in G2 and G3. Mice in G3 showed a more stable relapse rate than those in G1 or G2, but still maintained the same tendency for a rapid initial relapse followed by a gradual slower relapse (Fig. 2F).

### Effect of retention on the number of TRAP-positive cells during orthodontic relapse

TRAP staining was performed on tissue sections from the distobuccal root of the upper left first molar on r0 and r15 for mice in all three groups. At r0, there were much fewer osteoclasts on the mesial side in G3 (1.25 ± 0.82 cells/section) than in G2 (7.15 ± 1.24 cells/section) or G1 (12.4 ± 2.39 cells/section). There was a similar trend for osteoclast number at r15, with lower numbers found for all three groups as compared with those numbers at r0, respectively. These results suggest that fewer osteoclasts were activated after orthodontic tooth movement in mice with better retention (Fig. 3A and 3B). On the distal side, there was no significant difference among the three groups at r0 (G1, 0.35 ± 0.57 cells/section; G2, 0.25 ± 1.55 cells/section; G3, 0 ± 0 cells/section). By r15, osteoclast number had significantly increased in all groups as compared with the values at r0, but there was no significant difference among the groups (G1, 4.38 ± 1.09 cells/section; G2, 5.25 ± 0.68 cells/section; G3, 5.8 ± 0.78 cells/section) (Fig. 3A and 3C).

**Fig. 3.**
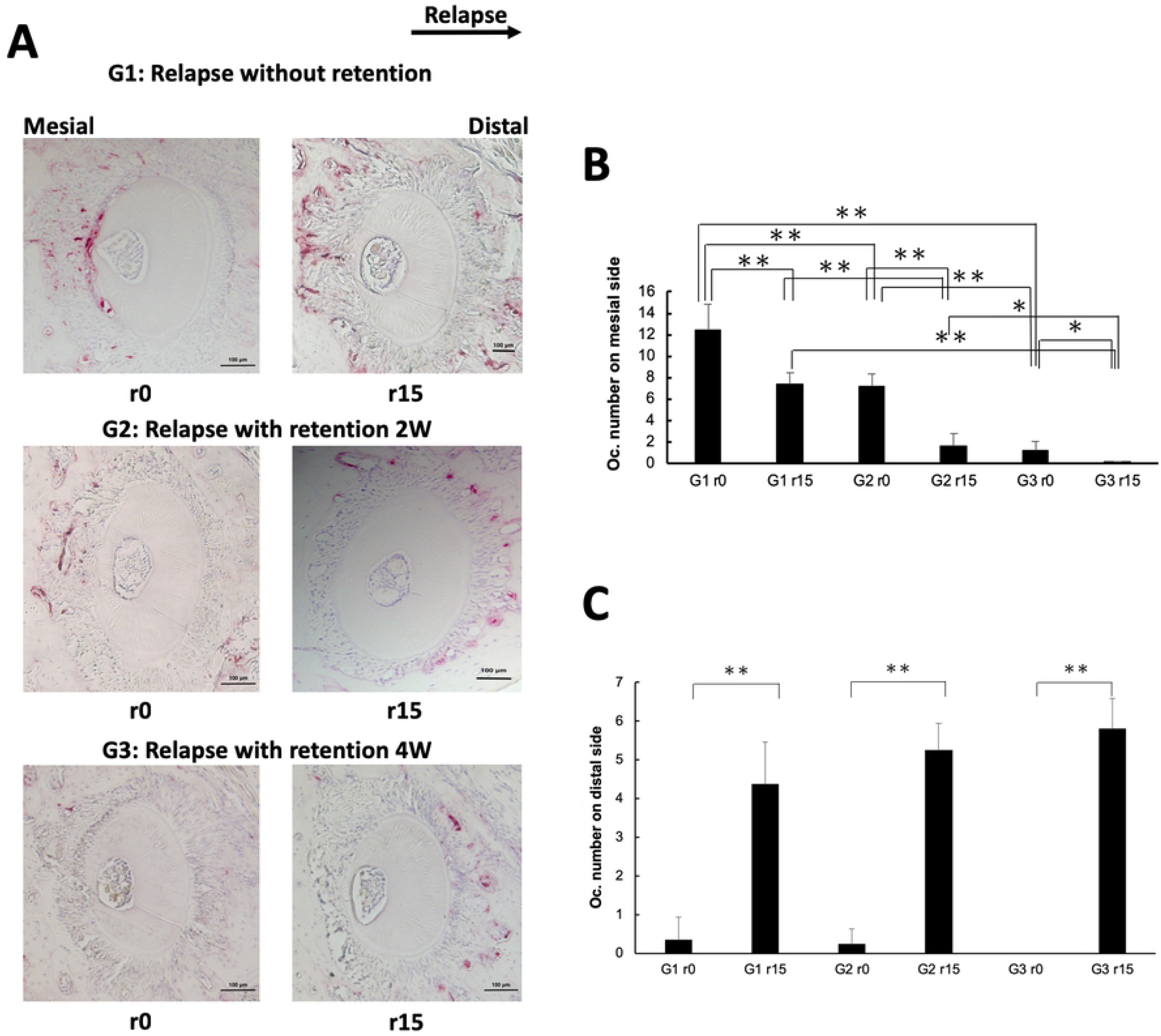
Evaluation of osteoclast number in horizontal sections of alveolar bone at the maxillary left first molar area. (A) Tartrate-resistant acid phosphatase (TRAP)-stained histological sections of the distobuccal root of the maxillary left first molar on relapse days 0 and 15 (r0 and r15) in each group. Arrow (top) indicates the direction of orthodontic relapse. Scale bars = 100 μm. (B) The number of TRAP-positive multinucleated cells on the mesial side of the distobuccal upper-left first molar were counted. Fewer osteoclasts were observed in mice treated with a longer retention period. *n =* 4 for each group. \**P* < 0.05; \*\**P* < 0.01. (C) The number of TRAP-positive multinucleated cells on the distal side of the distobuccal upper-left first molar. TRAP-positive cells were significantly increased at r15 in each experimental group as compared with numbers at r0. *n =* 4 for each group. \*\**P* < 0.01.

### Anti-c-Fms antibody inhibits orthodontic relapse by inhibiting osteoclastogenesis

The experimental design to examine the effect of inhibiting osteoclastogenesis is shown in the Figure 4A. Mice were treated with OTM for 12 days followed by 4 weeks of retention. Mice were then injected with anti-c-Fms or an equal volume of PBS to compare the effect of inhibiting osteoclastogenesis on relapse rates, as measured by changes in the space between M1 and M2. On r15, both groups showed a decreasing trend in orthodontic relapse as compared with r0. The space between M1 and M2 in the antibody group (124.06 ± 4.31 μm) was significantly greater than the space in the PBS (87.21 ± 6.59 μm) group (Fig. 4B) at r15. These findings suggest that orthodontic relapse was suppressed by injection with the anti-c-Fms antibody (Fig. 4C). To further confirm these findings, we calculated osteoclast number using histological analysis. On r15, more TRAP-positive cells were observed on the distal side of the alveolar bone in mice administered with PBS (6.8 ± 1.55 cells/section) as compared with those injected with anti-c-Fms antibody (1.45 ± 0.93 cells/section) (Fig. 4D and 4E).

**Fig. 4.**
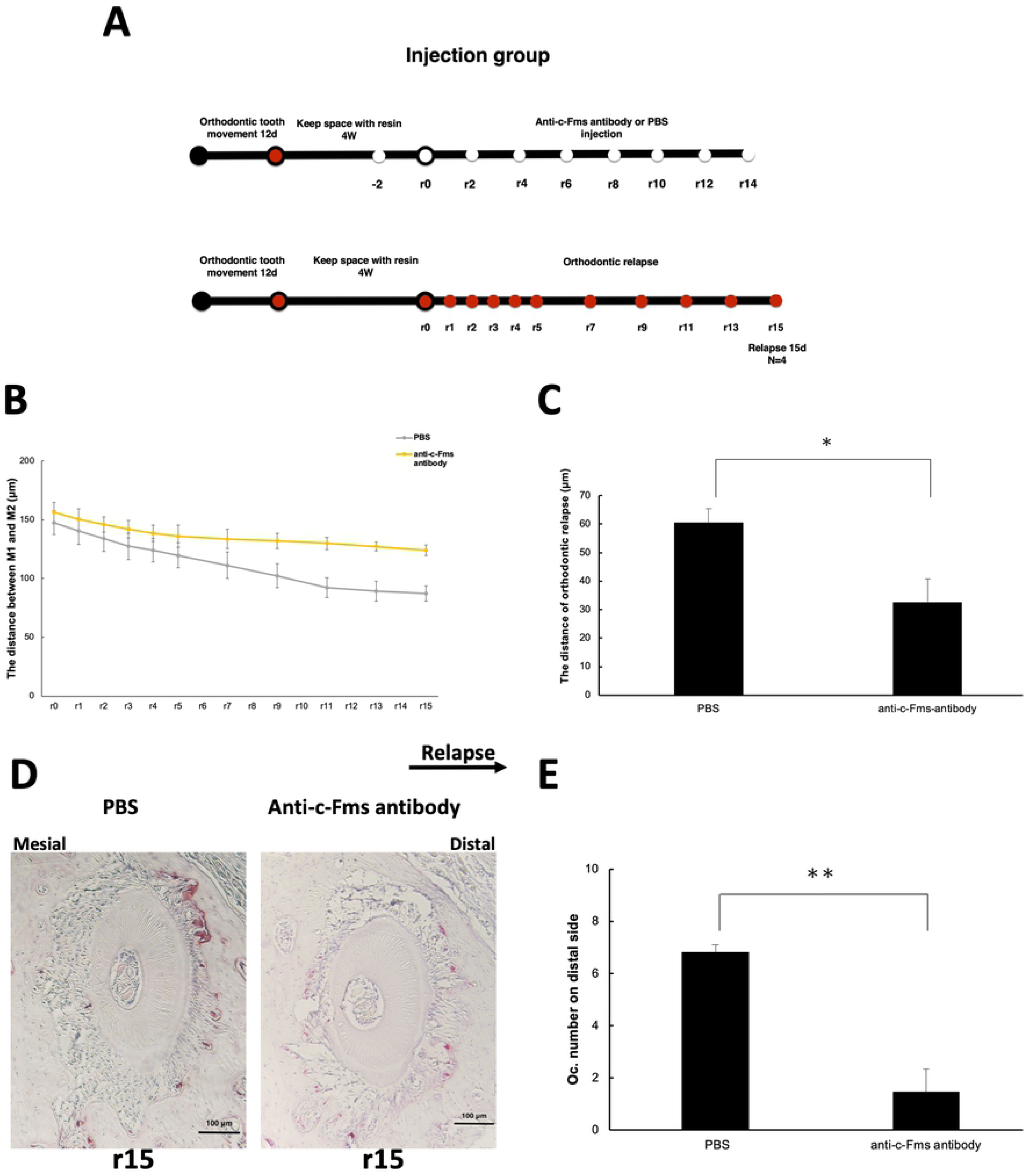
Evaluation of relapse distance and osteoclast activity after retention following injection with anti-c-Fms antibody. (A) Scheme for the experiment. Mice were treated with experimental loading for 12 days, followed by resin retention for 4 weeks. Mice were then injected with PBS or anti-c-Fms antibody every second day (as indicated by the white markers). Orthodontic relapse was measured on the days indicated by the red markers. (B) Changes in the amount of orthodontic relapse in each group. *n =* 4 for each group. (C) Comparison of relapse between mice treated with PBS or anti-c-Fms antibody on r15. *n* = 4 for each group. \**P* < 0.05. (D) Tartrate-resistant acid phosphatase (TRAP)-stained histological sections of the distobuccal root of the maxillary left first molar after treatment with PBS or anti-c-Fms antibody. r15, day 15 of relapse. Arrow shows the direction of orthodontic relapse. Scale bars = 100 μm. (E) The number of TRAP-positive multinuclear cells in mice treated with PBS or anti-c-Fms antibody.

## Discussion

Retention is one of the most important methods to prevent orthodontic relapse after orthodontic treatment. Despite this, the mechanism of orthodontic relapse remains unknown, with relapse often observed even after the effective use of a retainer in some patients. In this study, for the first time, we established a mouse model of orthodontic retention. The results show that mice treated with retention have a significantly shorter amount and more gradual rate of relapse when compared with mice not treated with retention. In addition, we found that a longer retention period lowered the relapse distance and rate of relapse. We also showed that orthodontic relapse was associated with the presence of osteoclasts, with fewer osteoclasts found in mice treated with a longer period of retention. Finally, we found that the anti-c-Fms antibody can inhibit orthodontic relapse by blocking osteoclastogenesis.

Intriguingly, all of the teeth subjected to forced orthodontic movement showed some evidence of tooth movement back toward their original positions. Faster and more significant movement was found following the removal of the coiled spring appliance (G1) than if the mice were subjected to some form of retention using light-cured resin (G2, G3). This finding indicates that an effective retention can help to reduce the degree of relapse post-treatment, with a longer retention period associated with a better outcome. However, it seems that retention cannot prevent relapse completely. We also found that bone remodeling might occur around the root during the retention period, with a longer retention period potentially associated with more new bone formation and less bone resorption.

Yet how orthodontic relapse occurs after retention is still unclear. We, therefore, analyzed osteoclast activity on histological sections of the distal buccal root of the upper left first molar on both the mesial and distal sides. On the mesial side, there was a high number of osteoclasts in mice not treated with retention (G1) and this number decreased in mice treated with retention, with longer retention associated with fewer osteoclasts. These findings indicate that bone resorption by osteoclasts still occurs during the period of retention on the mesial side. However, on the distal side, minimal osteoclast activity was detected at r0, but this increased proportionally in each group by r15. Overall, we surmise that osteoclast activity is a potential cause of orthodontic relapse.

In preliminary tests, we examined three different materials for fixation of the retention apparatus after OTM: a self-curing resin, a flowable composite, and light-cured resin. The self-curing resin and the flowable composite were difficult to control, even though the flowable composite was easy to initially position. In contrast, the light-cured resin showed good fixation and was thus chosen for subsequent experiments. The interdental space was maintained well by the light-cured resin with the use of a dental etching agent and a dental bonding agent.

Relapse occurred in all three groups following the removal of coil spring appliance (G1) or removal of the light-cured resin (G2, G3). The relapse rate was rapid initially but decreased gradually, and this is consistent with the findings in other studies [23-25]. Yoshida et al. found that, after 21 days of OTM, the space decreased from 526 μm to 108 μm on day 5 of the relapse period, and 71 μm on day 10, with a relapse rate of 83.6 μm/d and 7.4 μm/d, respectively [23]. In a rat model, Franzen et al. found that the relapse rate peaked on day 1, and then decreased gradually until study endpoint [25]. The relapse rate in our experiment showed a similar trend. Yadav et al. found that the space between M1 and M2 was 30.8 μm after 7 days of relapse in the mouse [26]. This finding corresponded well with our results of a mean movement of 24.62 μm among the mice not treated with retention.

OTM is achieved through repeated alveolar bone resorption by osteoclasts on the compressed side and stimulation of new bone formation by osteoblasts on the tension side. Osteoclasts were observed on the distal side at r15 in all of the groups, with numbers increasing as compared with values at r0. This finding suggests the involvement of osteoclasts during relapse. Yoshida et al. found prominent osteoclast-induced alveolar bone resorption during orthodontic relapse, and pointed to alveolar bone remodeling as a major reason for relapse in rats [23]. Our findings support these suppositions. In the current study, the distal side of the M1 tooth was subjected to tension during the force-triggered OTM. We speculate that there was still new bone formation during the retention period. One group reported new bone formation during remodeling of the alveolar bone at a rate of 6.7 μm/d in rat [27]. Given the lower rate of relapse and the smaller distal molar movement, we assume that mice subjected to retention may have increased bone mass on the distal side, which helps to resist relapse when the resin was removed.

It has been reported that stress stored within the periodontal and transeptal fiber system during OTM are released after removal of the orthodontic appliance [3], and that the periodontal ligament (PDL) can restore the original tooth arrangement during relapse [23]. Feng et al. identified the importance of PDL collagen recovery in early relapse [28]. Franzen et al. showed that the PDL width on the tension side increases following the application of orthodontic forces but becomes narrower at the end of the relapse period [25]. Considering all these findings, together with the present results, we suggest that retention increases the space between the first molar and the alveolar bone on the mesial side but decreases the space on distal side. The relapse force may also be lower because of a decrease in PDL pressure for the retention groups. Future studies are required to evaluate the change in PDL structure during relapse.

Physiological distal drift of mouse molars has previously been reported [29-31]. We found no significant difference in the relapse rate at r15 for any of the groups, with values on slightly more than 0. Although specific data related to distal drift are still unknown, in our present results, we found degrees of relapse tendency at 15 days after removal of the coiled spring appliance (G1) or removal of the light-cured resin (G2, G3). We surmise that the first molars most likely restored their physiological processes. Although humans have more complex physiology than mice, our mouse model showed a similar relapse trend as that seen in humans, and, although the space was retained successfully, there was still some evidence of orthodontic relapse. G3 showed the smallest relapse at r15.

We speculated that osteoclast-dependent orthodontic relapse would still occur after retention for 4 weeks. Therefore, we evaluated the effect of the anti-c-Fms antibody on orthodontic relapse after retention for 4 weeks. We previously reported that the anti-c-Fms antibody can inhibit osteoclastogenesis to decrease movement of the tooth back to its original position after forced repositioning [6]. Both orthodontic relapse and OTM have a similar pattern of cellular activity, with an increase in osteoclast activity on the compression side and a decrease on the tension side in rats [25]; we found similar patterns of cellular activity in our mouse model. Treatment with anti-c-Fms antibody reduced the amount of relapse and significantly decreased osteoclast number, as compared with that in the PBS group, suggesting that M-CSF and/or its receptor may be potential therapeutic targets for preventing orthodontic relapse after retention.

## Conclusion

In conclusion, we find that the amount and rate of relapse are shorter with a longer period of retention. We also show that orthodontic relapse is dependent on osteoclast number, which is high on the mesial side without retention, and reduces proportional to the length of the retention period. Anti-c-Fms antibody can inhibit osteoclastogenesis and, in turn, help to inhibit orthodontic relapse by reducing osteoclast activity. Thus, M-CSF and its receptor might be useful targets to inhibit osteoclast-mediated orthodontic relapse.

## Acknowledgments

This work was supported in part by a JSPS KAKENHI grant from the Japan Society for the Promotion of Science (No. 16K11776 to H.K.; No. 18K09862 to I.M.).

## Author Contributions

Jiawei Qi contributed to the conception, design, data acquisition, data analysis, data interpretation, and drafting of the manuscript. Hideki Kitaura contributed to conception, design, data acquisition, data analysis, data interpretation, and drafting and critical revision of the manuscript. Akiko Kishikawa, Saika Ogawa, Wei-Ren Shen, Fumitoshi Ohori, Takahiro Noguch, Aseel Marahleh, Yasuhiko Nara, and Itaru Mizoguchi contributed to data acquisition and analysis. All authors provided final approval and agree to be accountable for all aspects of the work.

